# Evidence for a functional interaction between the respiratory syncytial virus fusion and attachment proteins in the envelope of infectious virus particles

**DOI:** 10.1101/2022.12.18.520517

**Authors:** Tra Nguyen Huong, Boon Huan Tan, Richard J. Sugrue

## Abstract

We have examined the interaction between the respiratory syncytial virus (RSV) F and G proteins on the surface of infected cells during multiple cycle infection using a low multiplicity of infection (MOI) model, and on the surface of virus particles that were isolated from infected cells. A combination of the proximity ligation assay (PLA) and confocal microscopy was used to demonstrate the interaction between the F and G proteins within the virus filaments on infected cells. Co-precipitation of the F and G proteins was confirmed using detergent extracts prepared from infected cells and in detergent extracts prepared from purified virus particles. The influence of the G protein in mediating virus spread in the low MOI model was further examined using the recombinant virus isolates rg224RSV (that expresses all virus proteins) and rg224RSV-ΔG (which does not express the G protein). While cells could be initially infected by both viruses, the rg224RSV-ΔG virus exhibited severely impaired localised virus transmission in the multiple cycle infection assay. Collectively these data provide evidence that the F and G proteins interact within the envelope of RSV particles, and suggests that this interaction may promote virus transmission. The interaction between these proteins in a single protein complex represents a potential new target for the development of antivirus strategies and in the development of RSV vaccine candidates.

## Introduction

The mature Respiratory Syncytial Virus (RSV) particle assembles at the apical cell surface as cell-associated filamentous structures [1], and during RSV infection it is the virus filaments that mediate virus transmission [2, 3]. The virus filaments are bounded by a virus envelope in which the G protein and virus fusion (F) protein are inserted. The G protein is the RSV cell-attachment protein [4] and it undergoes extensive glycosylation to form the mature 90 kDa G protein [5]. The F protein initiates fusion of the virus envelope and cell membrane during virus entry into the cell. It is initially expressed as a single polypeptide chain (F0), and it is subsequently cleaved by furin into the F1 (50 kDa) and F2 (20 kDa) subunits [6–8]. Several models have been put forward to explain the mechanism of RSV-induced membrane fusion [9]. It is suggested that the mature F protein initially exists as a labile pre-fusion state, which is subsequently converted into a stable post-fusion molecule during membrane fusion [10, 11]. While the structural basis for the fusion process mediated by the RSV F protein is understood [9, 12], the cue that converts the labile pre-fusion state into the stable post-fusion form during virus infection is not established. In many paramyxoviruses the interaction of the attachment protein with the cell receptor mediates structural changes in the F protein that are required during membrane fusion [13]. In this context we have previously demonstrated that a protein complex involving the F and G protein forms on the surface of infected cells [14]. This earlier study relied on the co-immunoprecipitation of detergent extracts obtained from virus-infected cells. However, this earlier study did not specifically address if this protein complex was incorporated into the envelope of virus filaments (i.e. infectious virus particles). We have therefore extended our earlier findings to determine if this protein complex was formed in the envelope of infectious virus particles. The presence of this complex in virus particles would have important implications towards understanding how the process of F protein mediated cell entry is regulated. It could also potentially lead to both novel antivirus strategies that are based on destabilising this interaction and in the development of recombinant RSV subunit vaccines.

## Materials and Methods

### Virus and cell preparation

The rg224RSV-FSG and rg224RSV-FS viruses have been described previously [15, 16]. The RSV A2, rg224RSV-FSG and rg224RSV-FS viruses were prepared in HEp-2 cells as described previously [17, 18]. Infections were performed in HEp-2 cells at the desired multiplicity of infection in DMEM with 2% FCS in a humidified chamber at 33 °C with 5% CO_2_. The HEp2 cells were maintained in DMEM with 10% FCS incubated at 37 °C with 5% CO_2_.

### RSV multiple cycle virus infection assay

The cell-associated and cell-free (released) virus infectivity was measured as described previously [19]. Briefly, HEp2 cells that were 95% confluent were infected with RSV using a multiplicity of infection of between 0.001 and 0.0001 as indicated. At specific time intervals the cell-free virus infectivity was recovered by harvesting the tissue culture supernatant. The cell-associated virus was recovered by scrapping the cells into the same volume of pre-warmed DMEM (2% FCS), and the virus infectivity released from the cells by two cycles of rapid freeze thaw. Contaminating cellular material was removed from each virus preparation by centrifugation at 5000g for 10 minutes at 4°C. The recovered viral infectivity in each fraction was assessed on HEp-2 cells by microplaque assay.

### Microplaque titration of RSV infectivity

The recovered viral infectivity was assessed on HEp-2 cells using a microplaque assay as described previously [20] but with minor modifications. Briefly, at 48 hrs post-infection (hpi) the HEp-2 cell monolayers were incubated with an anti-RSV (Novacastra) and anti-mouse IgG conjugated to Alexa 488. The stained microplaques were counted using a Nikon Eclipse 80i (Japan) fluorescent microscope.

### Antibodies and specific reagents

The anti-mouse and anti-rabbit IgG conjugated to Alexa 488 and Alexa555 (Molecular Probes), anti-NP (Chemicon, San Diago, CA, USA) and Alexa Fluor™ 488 Phalloidin (ThermoFisher) were purchased. The RSV M2-1 antibody has been described previously [21–23], and the F protein polyclonal and anti-G were gifts from Jose Melero (Madrid, Spain) and Geraldine Taylor (IAH, UK) respectively.

### Proximity Ligation assay

The cells were fixed using 4% PFA in PBS and then incubated with anti-G and anti-F antibodies (either in combination or singularly as required). The proximity Ligation Assay (PLA) was then performed using the Duolink^®^ In Situ Detection Red as recommended by the manufacturer (Sigma Aldrich). After the PLA reaction was performed, the cells were then incubated with the anti-G or anti-F antibodies and the antimouse or anti-rabbit IgG conjugated to Alexa 488 as appropriate. Cells were washed with PBS and mounted on glass slide using mounting media (Dakocytomation Fluorescence Mounting Media, Dako, USA). The cells were examined using either a Nikon Eclipse 80i (Japan) fluorescent microscope or by using a Zeiss 710 confocal microscope with Zeiss software.

### Immunofluorescence microscopy

This was performed as described previously [24]. Briefly, the cells were fixed using 4% (w/v) paraformaldehyde in PBS at 4°C for 20 mins and washed using PBS at 4°C. For Alexa Fluor™ 488 Phalloidin staining the cells were fixed using 3% (w/v) paraformaldehyde in PBS. The cells were permeabilised using 0.1% (v/v) triton X100 in PBS at 4°C for 15 mins. The cells were stained using the primary and secondary antibody combinations (Alexa 488 or Alexa 555 as appropriate). Alexa Fluor™ 488 Phalloidin staining was performed using the manufacturer’s instructions. The cells were examined using either a Nikon Eclipse 80i (Japan) fluorescent microscope or by using a Zeiss 710 confocal microscope with Zeiss software as appropriate.

### Immunoblotting analysis

The cells were washed twice using sterile PBS (at 4°C) and then extracted directly into Boiling Mix (1%(w/v) SDS, 5% mercaptoethanol (v/v) in 20mM Tris/HCL, pH 7.5). The extracted cells were immediately heated at 100°C for 2 min and then clarified by centrifugation (13,000 x *g* for 2 min). The proteins were separated by SDS-PAGE and transferred by Western blotting onto nitrocellulose membranes. Protein bands were then probed with the relevant primary antibody and corresponding secondary antibody conjugated to HRP, and visualized using the ECL detection system (GE Healthcare).

### Surface protein biotinylation

This was performed with 0.5 mg/ml EZ-Link Sulfo-NHS-LC-LCBiotin (Pierce Biotechnology) solution in PBS pH 8 as described previously [14]. The cell monolayers were treated with 0.5 mg/ml EZ-Link Sulfo-NHS-LC-LCBiotin (Pierce Biotechnology) solution in PBS pH 8 for 1 h at 20°C. The cells were washed using PBS with 2mM lysine and then the cells were lysed using ice-cold extraction buffer (PBS pH 8, 1% Nonidet P-40 (BDH), 1 mM EDTA, 2mM lysine and 2 mM PMSF) at 4°C for 20 min and clarified by centrifugation (13,000g, 10min) at 4°C. The biotinylated proteins were isolated by immunoprecipitation and transferred onto PVDF membranes by western blotting and detected using streptavidin-HRP. Protein bands were quantified by using ImageJ (ver IJ1.46r) to analyse the protein bands on autoradiographs. Protein profiles were obtained or protein bands to be quantified were delineated, and the relative intensities determined and compared with the background intensity in control lanes.

### Biotinylation of RSV particles

HEp2 cells were infected with RSV using a multiplicity of infection of 0.05 and at 60 hpi the infectious virus particles were isolated using ultracentrifugation as described previously [18]. The virus particles were suspended in PBS pH 8 with 0.5 mg/ml EZ-Link Sulfo-NHS-LC-LCBiotin for 1 h at 20°C. The reaction was quenched by adding 20mM lysine and after 10 mins the virus particles recovered by centrifugation (284,000 × g) for 90 mins at 4°C. The virus particles were suspended in Hanks buffered saline solution HBSS with 20mM lysine, and extracted using an equal volume of 2% (v/v) Nonidet P-40 in HBSS with 20mM lysine at 4°C for 20 mins. The detergent extract was clarified by centrifugation (13,000g, 10min) at 4°C and the proteins isolated by immunoprecipitation [14].

## Results and discussion

We have previously described a low multiplicity of infection (MOI) model for examining the multiple cycle infection in RSV-infected cell monolayers [19, 25]. Under these conditions localised virus transmission in cell monolayers was observed when using a moi of between 0.001 and 0.0001. This phenomenon is a robust feature of all permissive cell lines infected with RSV. High mutation rates that are associated with RNA viruses can lead to the formation of defective interfering (DI) particles that are potentially non-infectious or that exhibit impaired infectivity [26]. The presence of DI particles in virus preparations have also been shown to significantly influence the virus transmission processes in permissive cells [27–29], and unless otherwise specified, in this current study the HEp2 cells were infected with RSV using a MOI of 0.0001. Under these conditions the DI particles that would be expected to be present in the initial virus inoculum would be sufficiently diluted out and individual cells in the monolayer would only be infected with individual and fully infectious virus particle. This allowed both RSV transmission in the cell monolayer and the interaction between the F and G proteins to be examined with the minimal involvement of the DI particles. HEp2 cells were infected with RSV and at between 1 and 3 days-post-infection (dpi) the cell monolayers were co-stained using anti-G and Alexa Fluor™ 488 Phalloidin and imaged using immunofluorescence (IF) microscopy (Fig. 1A). While anti-G staining allowed infected cells to be visualised, the Alexa Fluor™ 488 Phalloidin stained the F-actin and confirmed the integrity of the cell monolayer. At 1 dpi mainly singly infected cells were detected within the cell monolayer, while at 2dpi larger infected cell clusters that consisted of 25±15 cells per cluster was detected, and the anti-G staining was restricted to these infected cell clusters. At 3 dpi enlarged cell clusters were detected, and their appearance correlated with the start of syncytia formation. At each time of infection the cell-associated (CA) and cell-free (CF) virus infectivity was separated and quantified by plaque assay (Fig. 1B). At between 1 and 3 dpi a continuous increase in the CA-virus infectivity was detected, while the appearance of CF-virus infectivity was only detected by 3 dpi. This pattern of virus spread confirmed our previous observations demonstrating the localised cell-to-cell transmission of RSV in cell monolayers [19].

**Figure 1.**
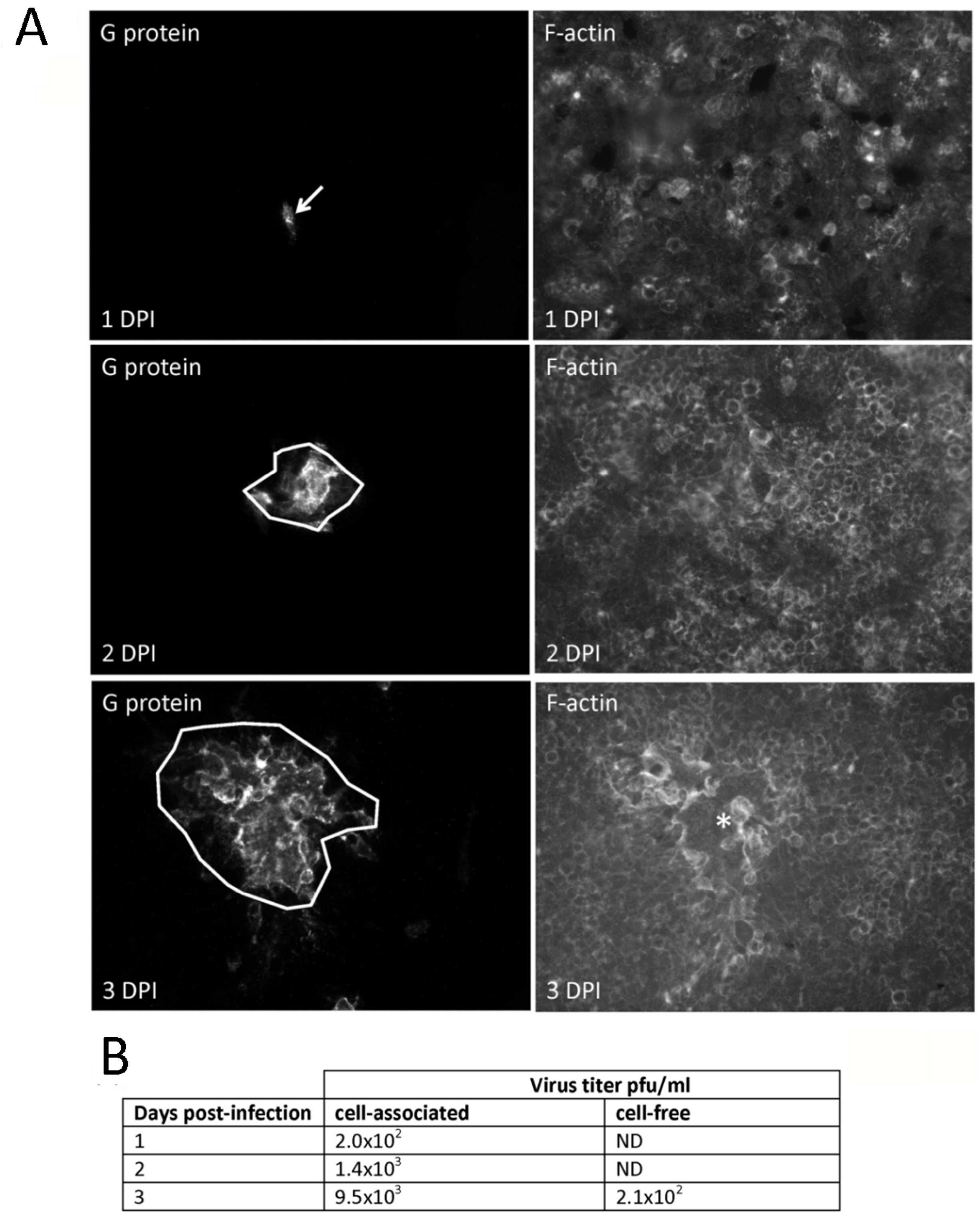
Distribution of the F and G proteins on cells infected using the low multiplicity of infection model. HEp2 cell monolayers were RSV-infected using a multiplicity of infection (MOI) of 0.0001 and at between 1 and 3 days post-infection (dpi). **(A)** The monolayers in the tissue culture dish were fixed using 3% (w/v) paraformaldehyde and stained using anti-G followed by anti-mouse IgG conjugated to Alexa 555 and Alexa Fluor™ 488 Phalloidin. The stained cells were then viewed using fluorescence microscopy (objective x40 magnification). The infected cell clusters (white arrow) and syncytial formation (*) in the monolayer are highlighted, and Alexa Fluor™ 488 Phalloidin staining (F-actin) of the monolayer shows the cell monolayer. **(B)** The cell-associated and cell-free virus infectivity was recovered at each time of infection as described previously [19] and the infectivity assessed by microplaque assay. ND means not detected.

At 2 dpi the HEp2 cells were also co-stained using anti-F and anti-G and the infected cell clusters examined by IF microscopy (Fig. 2 A-C), which showed strong co-staining within the virus filaments. Examination of the co-staining pattern in more detail indicated that while anti-G staining was observed along the whole length of the virus filaments, increased anti-F staining at the distal end of the virus filaments suggested the increased concentration of the F protein at the ends of the virus filaments (Fig. 2 B and C). These imaging data suggest a compartmentalisation of the F protein distribution in the virus filaments, and this is similar to the observation in RSV VLPs that are formed by the co-expression of the F and G proteins [30]. At 2 dpi the HEp2 cells were also examined by confocal microscopy (Fig. 3A), which also indicated strong anti-G and anti-F co-staining within the virus filaments at the distal ends of the virus filaments. Examination of the pixel distribution at the periphery (Fig. 3B (i) and (ii))) and top (Fig. 3C (i) and (ii)) of the infected cell clusters confirmed high levels of co-localisation between the anti-F and anti-G antibody staining within virus filaments (Pearson coefficient = 0.78±0.04).

**Figure 2.**
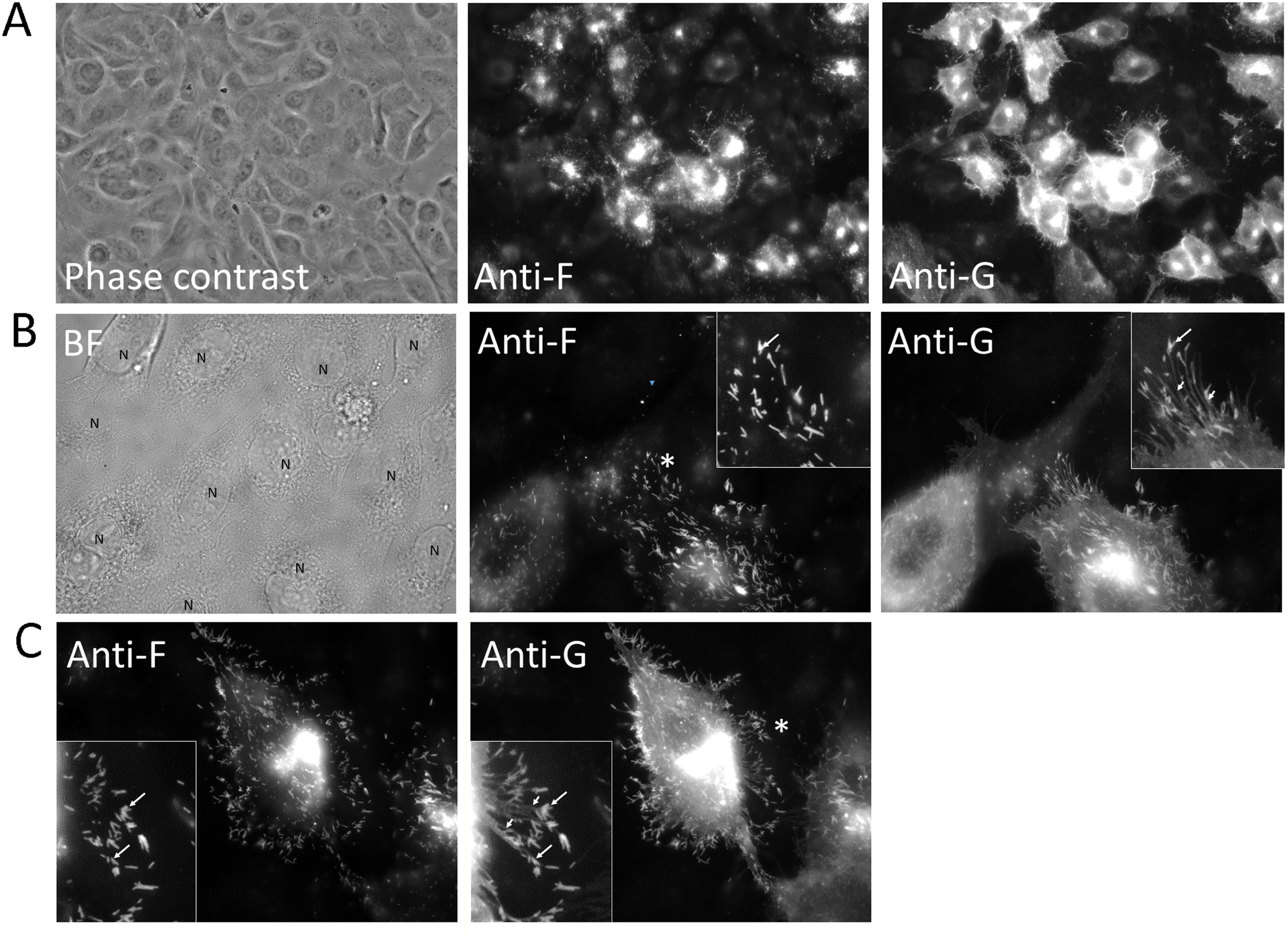
Distribution of the F and G proteins on cells infected using the low multiplicity of infection model. HEp2 cell monolayers were RSV-infected using a MOI of 0.0001 and at 2 dpi the cells were co-stained with anti-F and anti-G. **(A)** the stained cells were examined using phase contrast microscopy to view the cells in the field of view and by immunofluorescence (IF) microscopy to visualise the anti-G and anti-F staining on infected cells (objective x40 magnification). **(B and C)** As indicated, the stained cells were examined using bright-field (BF) microscopy to view the cells in the field of view and using IF microscopy to visualise the anti-G and anti-F staining on representative infected cells (objective x100 (oil) magnification). In the BF microscopy images the nuclei (N) of cells in the field of view are highlighted. In the IF microscopy images the insets are enlarged images taken from the cell at the region indicated (*). The anti-G stained (short white arrows) and anti-G and anti-F co-strained (long white arrows) regions of the virus filaments are also indicated in the insets.

**Figure 3.**
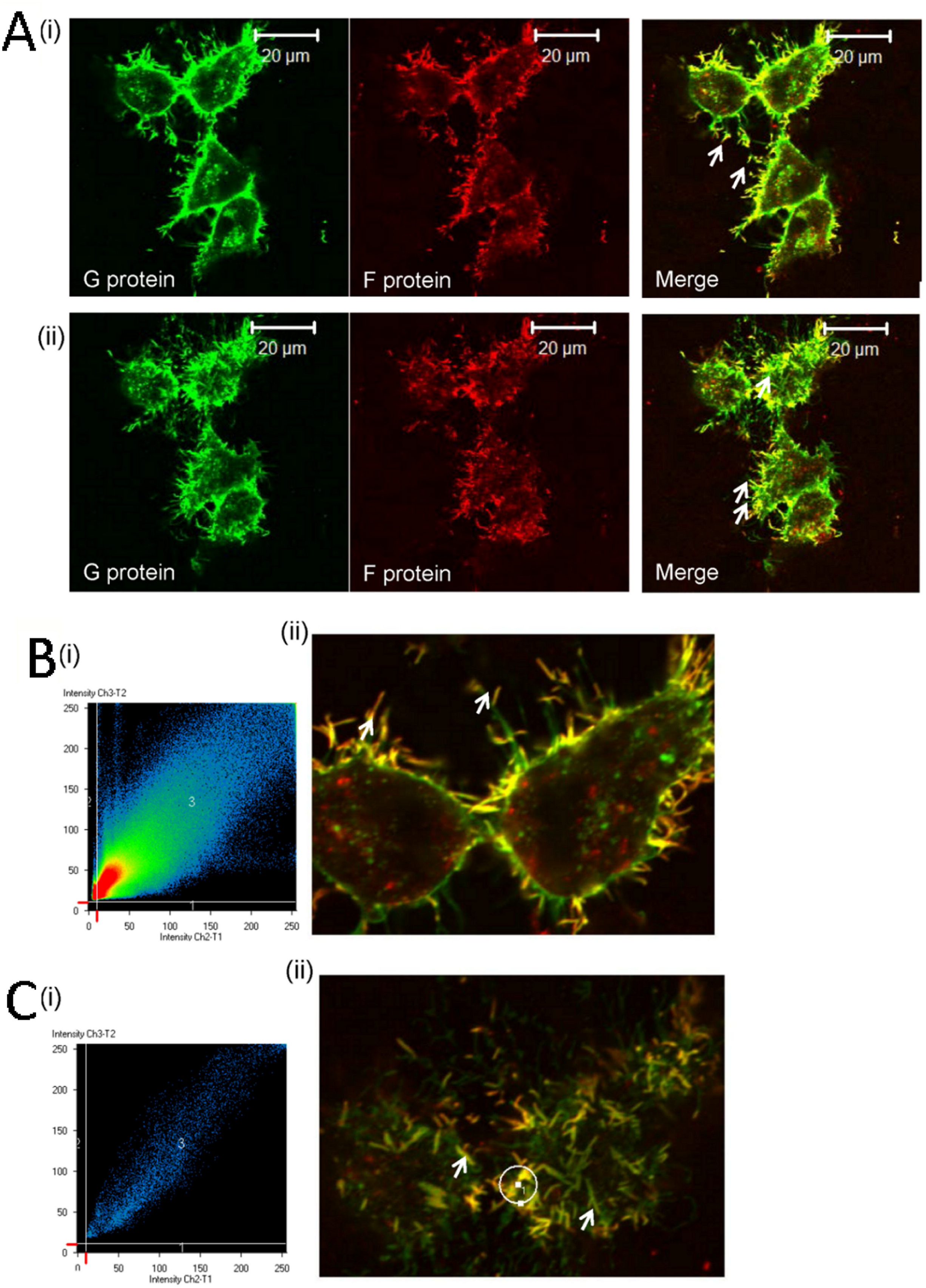
Distribution of the F and G proteins on cells infected using the low multiplicity of infection model. HEp2 cell monolayers were RSV-infected using a MOI of 0.0001 and at 2 dpi the cells were co-stained with anti-F and anti-G, and **(A)** the infected cell cluster examined by confocal microscopy at an optical plane that allows visualisation of the (i) cell periphery and (ii) cell top. The virus filaments are highlighted (white arrows). **(B and C)** Colocalisation analysis was performed by examining the pixel distribution in the co-stained cells at a focal plane that allows the **(B)** the cell periphery and **(C)** the cell top to be examined. **B**(i) shows a scatter plot of the entire image in **B**(ii), while **C**(i) is a scatter plot of the area highlighted in **C**(ii) by white circle. The virus filaments are highlighted (white arrows).

The high level of co-localisation was suggestive of an interaction *in situ*, and this possibility was examined further using the proximity ligation assay (PLA), which is an immuno-PCR based-assay that allows detection of interacting proteins *in situ* [31]. Although this assay cannot quantitate the number of interactions involving specific proteins, it is able to provide a qualitative indication of interacting proteins in intact cells. In the PLA the cells were incubated with anti-F and anti-G antibodies, followed by secondary antibodies that are conjugated to complementary DNA sequences that form a PCR primer pair. When the antibodies are in close proximity, the reaction products appear as distinct bright red spots when viewed by IF microscopy. HEp2 cells were either mock-infected or infected with RSV, and at 2 dpi the cells were incubated with anti-G and anti-F antibodies and the PLA reaction performed. The cells were subsequently labelled with anti-mouse IgG-Alexa 488 to detect the anti-G staining on the infected cell monolayer, which allowed us to examine the distribution of the virus filaments on the infected cells in the context of the PLA signal. The PLA signal was absent on the mock-infected cells (Fig. 4A), and on the infected cell monolayer, the PLA signal was restricted to the anti-G stained infected cell clusters (Fig. 4B (i) and (ii)). The PLA signal was absent on the negative control assays in which the PLA performed on infected cells that were incubated using either only anti-G or anti-F antibody (Fig. 4C (i) and (ii)). This demonstrated that the positive PLA reaction was not due to antibody cross reactivity. Collectively, these data indicated a positive PLA reaction in the virus-infected cells, and provided direct evidence for an interaction between the F and G proteins in the low MOI multiple cycle infection model.

**Figure 4.**
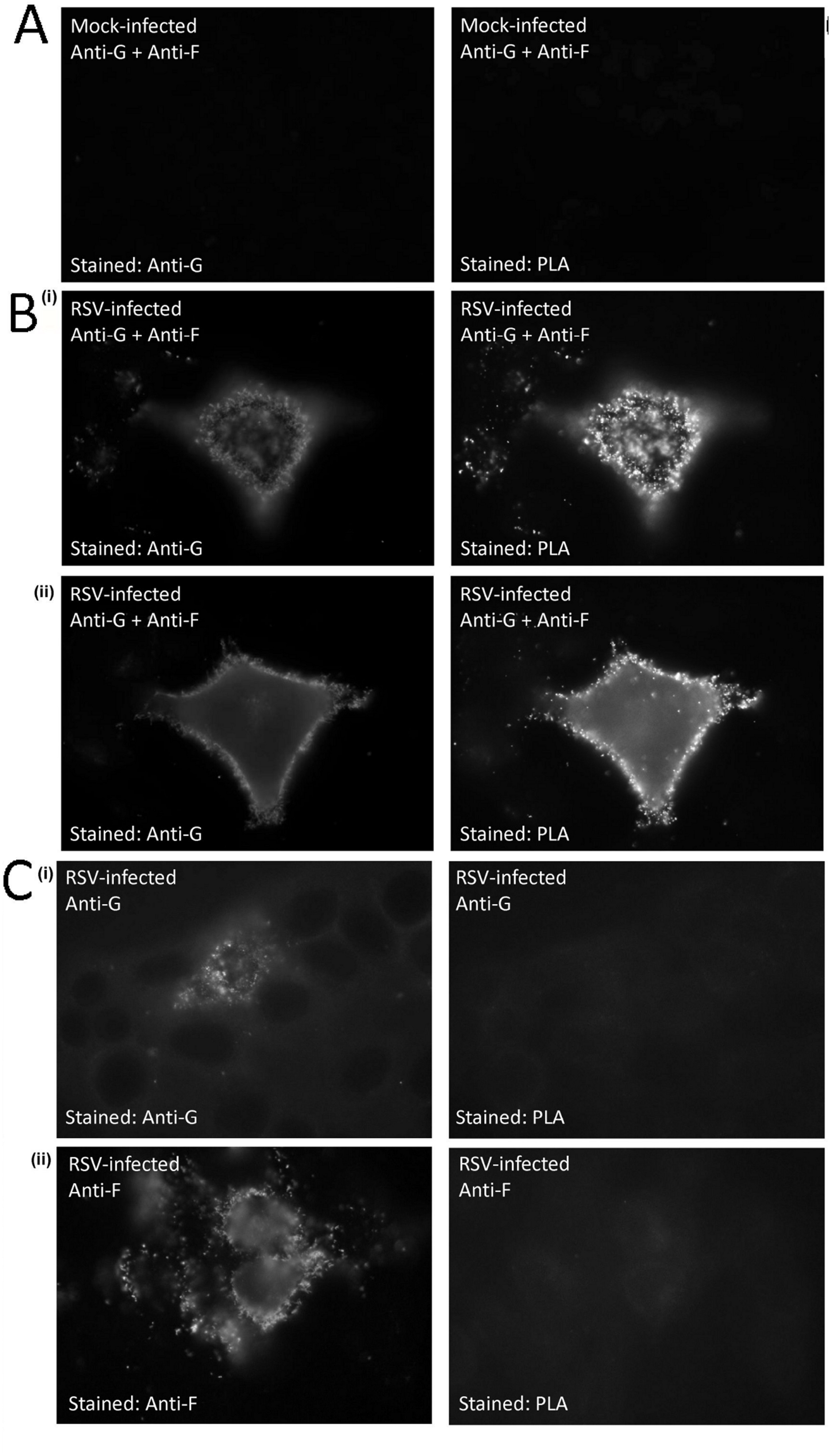
The F and G proteins interact on the surface of virus filaments. HEp2 cell monolayers were **(A)** mock-infected and **(B)** RSV-infected using a multiplicity of infection of 0.0001, and at 2 days-post infection (dpi) the cells were labelled with anti-F and anti-G and the proximity ligation assay (PLA) performed and the cells stained using anti-G. In each case the antibody staining and PLA signal is shown. (**B**(i)) highlights the cell top and B(ii) the cell periphery. **(C)** Cells were also singly labelled using either (i) anti-G or (ii) anti-F and the PLA performed. The cells were then stained with anti-G and anti-F as indicated.

The co-stained infected cells were examined in greater detail using confocal microscopy to determine the distribution of the PLA signal in the context of the anti-F stained virus filaments at the cell periphery (Fig. 5A(i)-(iii)). High levels of PLA staining on the anti-F stained virus filaments was visible and examined the pixel distribution of the PLA signal and anti-F staining (Fig. 5B(i) and (ii)) at regions of the images where the PLA signal was present. This revealed high levels of co-localisation between the PLA signal and the anti-G stained virus filaments (Pearson’s correlation coefficient = 0.68±0.05). Similarly, analysis of the antibody staining intensity profile across the virus filaments (Fig. 5C(i) and (ii)) where the PLA signal was present showed good correlation between the PLA signal and anti-F staining intensities. These analyses confirmed the presence of the PLA signal on the anti-F stained virus filaments, which accounted for greater than 95% of the total PLA signal detected indicating close association of the F and G proteins in these structures.

**Figure 5.**
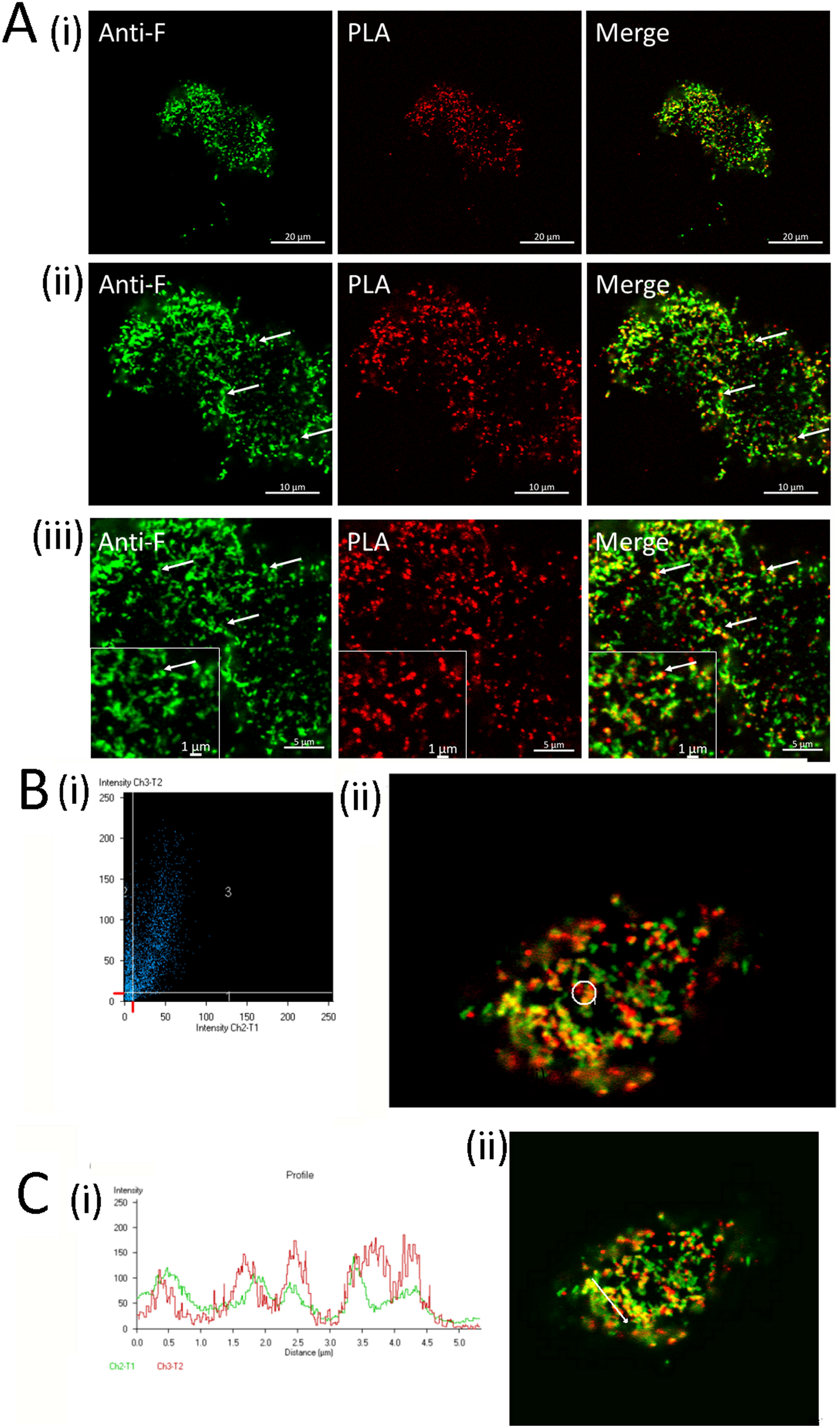
The PLA signal is localised to virus filaments. HEp2 cell monolayers were infected with RSV using a multiplicity of infection of 0.0001, and at 2 days-post infection (dpi) the cells were labelled with anti-F and anti-G and the proximity ligation assay (PLA) performed. **(A)** The cells were stained using anti-F, and the anti-F staining and PLA signal was examined by confocal microscopy at a focal plane that allowed the visualisation of the virus filaments at different magnifications ((i) to (iii)). Inset in (iii) is an enlarged image taken from each plate. The virus filaments are highlighted (white arrow). **(B)** Co-localisation analysis was performed by examining the pixel distribution in the anti-F stained cells following the PLA assay. (i) Is a scatter plot of the region highlighted in (ii). **(C)** Distribution of staining intensity in the anti-F stained cells following the PLA assay. (i) The intensity profile of the area demarcated by the white line in (ii).

We examined the interaction of the F and G proteins under low moi conditions using an established cell surface biotinylation assay that has been described previously [14]. Cells were either mock-infected or with RSV infected using a moi of 0.001, and at 2 dpi the cells were surface biotinylated and cell lysates prepared. Analysis of the total biotinylated proteins in the mock-infected and RSV-infected cell lysates revealed biotinylated proteins up to 200 kDa in size (Fig. 6A(i)). The cell lysates were immunoprecipitated using anti-F antibody which revealed the appearance of the F1 protein and a 90 kDa protein corresponding in size to the G protein in the RSV-infected cells lysates. Similarly the appearance of a 90 kDa G protein and lower levels of the 50 kDa F1 protein were observed by immunoprecipitation of the RSV-infected cell lysates with anti-G. These biotinylated protein species were not detected in lysates prepared from mock-infected cells, nor were they detected following immunoprecipitation with anti-NP (an antibody that recognises the influenza virus nucleoprotein). The ratio of the G to F1 protein bands was approximately 10:1 and 1:1 in the immunoprecipitation assay using anti-G and anti-F antibodies respectively (Fig. 6A(ii)).

**Fig. 6.**
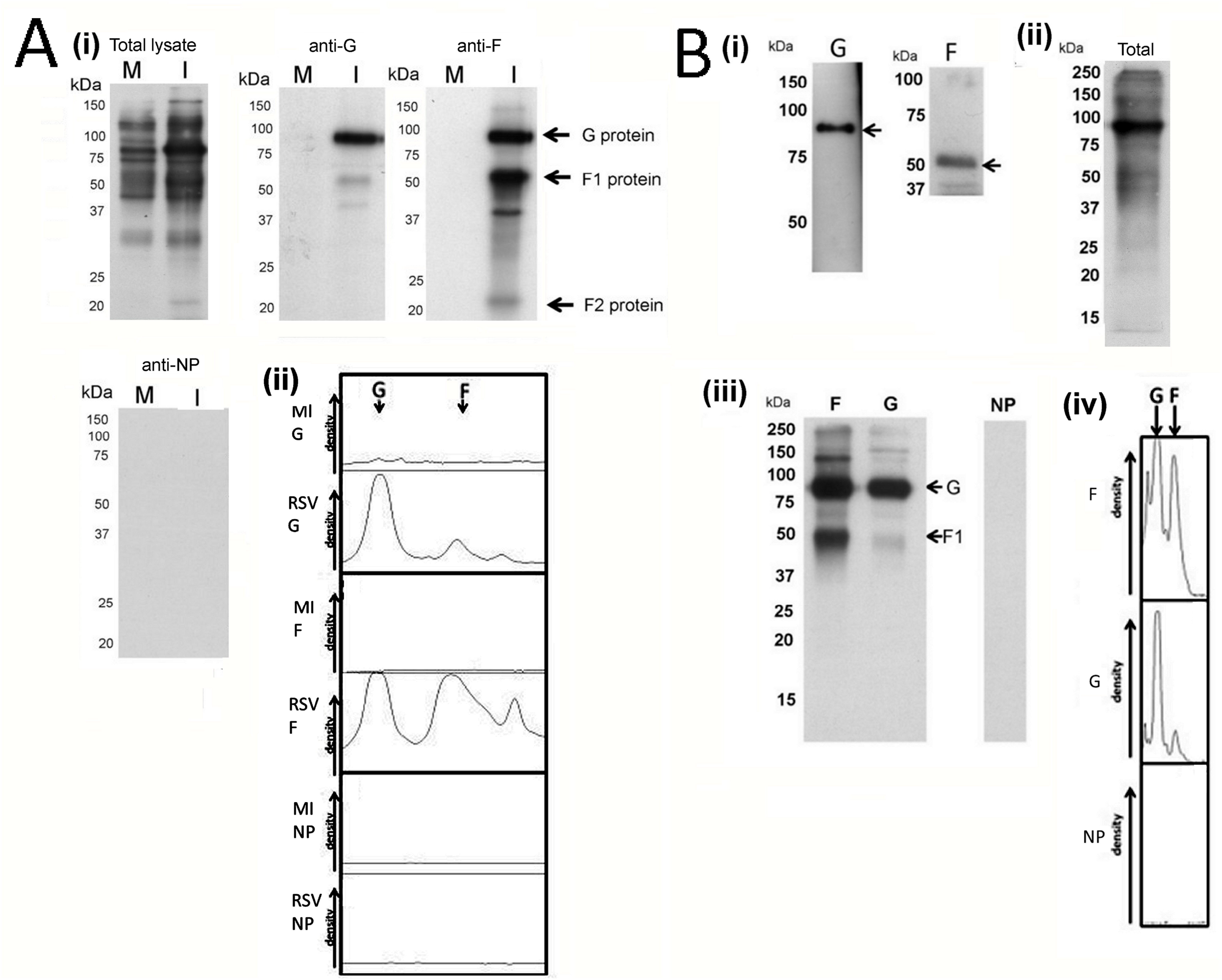
Co-precipitation of the F and G proteins in the low multiplicity of infection model and in purified virus particles. **(A)** HEp2 cell monolayers were mock-infected (M) and RSV-infected using a multiplicity of infection of 0.001 (I) and at 48 hr post-infection (hpi) the cells were surface-biotinylated. (i) the clarified detergent extracts were either examined directly (total lysates) or immunoprecipitated using anti-G, anti-F or anti-NP as indicated. Protein species corresponding in size to the F1, F2 and G proteins are also indicated. (ii) Densitometry analysis of the immunoprecipitation from mock-infected (MI) and virus-infected (RSV) cells are shown for the immunoprecipitation with anti-G (G), anti-F (F) and anti-NP (NP). **(B)** The virus particles were harvested as described previously [18] and (i) The virus preparation was examined by immunoblotting with anti-G (G) and anti-F (F), and protein bands corresponding in size to the mature G protein and the F1 protein are indicated. **(ii)** The virus preparation was surface-biotinylated and the proteins in the virus fraction was analysed by Western blotting. The total surface biotinylated proteins in the clarified detergent extract are shown. (iii) The surface biotinylated virus was detergent extracted and immunoprecipitated using anti-F (F), anti-G (G) and anti-NP (NP). Protein species corresponding in size to the F1 protein and the mature G protein are indicated. (iv) Densitometry analysis of the immunoprecipitation assays with anti-F (F), anti-G (G) and anti-NP (NP).

The interaction between the F and G proteins was also examined in virus particles that were isolated from HEp2 cells by ultracentrifugation as described previously [14]. Immunoblotting of the virus preparation with anti-F and anti-G showed proteins of the expected size for the 50 kDa F1 protein and the 90 kDa G protein respectively (Fig. 6B(i)). The virus preparation was surface-biotinylated and the analysis of the total-labelled proteins in the detergent virus extract revealed a prominent protein band at approximately 90 kDa, and lower levels of biotinylated protein species up to 250 kDa were also observed (Fig. 6B(ii)). Immunoprecipitation with anti-F showed the presence of the 50 kDa F1 protein together with a 90 kDa protein whose size was consistent with the G protein, while immunoprecipitation with anti-G showed a prominent 90 kDa G protein species, together with a minor protein band corresponding in size to the F1 subunit. (Fig. 6B(iii)). The absence of biotinylated protein species following immunoprecipitation with anti-NP (non-specific antibody) demonstrated the specificity of the immunoprecipitation assay. The ratio of the G to F1 protein bands was approximately 10:1 and 1:1 in the immunoprecipitation assay using anti-G and anti-F antibodies respectively (6B(iv)) and was similar to that observed in the biotinylation analysis on virus-infected cells described above.

A single protein complex involving the F and G proteins within the envelope of infectious RSV particles provided evidence for a functional linkage between the activity associated with the G protein (cell attachment) and the F protein (membrane fusion). The surface expressed virus glycoproteins were detected by biotinylation of surface accessible lysine residues and the F and G protein may be labelled to varying degrees (e.g. due to differences in the abundance and surface exposure of the lysine residues). It was therefore not possible to determine the stoichiometry of the F and G proteins in this protein complex. However, this analysis revealed that the relative amounts of the F and G proteins co-immunoprecipitated using either anti-F or anti-G differed. While an approximate ratio of 1:1 was noted by immunoprecipitation with anti-F, a lower level of the F1 protein was co-precipitated by immunoprecipitation with anti-G. The difference in the ratio in the band intensity observed with the different antibodies suggested that while the F and G protein interact on the surface of cells, not all the surface expressed G protein interacts with the F protein. These differences suggested that the F protein may interact with only a subpopulation of the total G protein that is present in the virus envelope. This conclusion is consistent with the imaging analysis which indicated increased levels of the F protein at the distal ends of the virus filaments, and that the co-localisation between anti-F and anti-G staining was mainly observed at the distal ends of the anti-G stained virus filaments.

In a final analysis we examined if the absence of the G protein impaired RSV transmission in the low MOI infection model. This was performed by comparing the transmission of wild-type RSV virus (rg224RSV) that expresses all three virus glycoproteins with a virus that does not express the G protein (rg224RSV-ΔG). Both viruses were engineered to express enhanced green fluorescent protein (EGFP), which is an additional marker to assess infection. The replication characteristics of both viruses have been previously extensively characterised, and the rg224RSV virus has been shown to exhibit similar replication characteristics to the non-EGFP expressing parent virus (e.g. [15]). HEp-2 cell monolayers were infected with rg224RSV or rg224RSV-ΔG, and at between 1 and 4 dpi the cell monolayers were stained using anti-F antibody and visualised using IF microscopy (Fig. 7A). In rg224RSV-infected cells small clusters of infected cells were detected at 2 dpi that increased in size by 3 dpi, and by 4 dpi syncytia formation had occurred. The pattern of spread of the rg224RSV virus in the low MOI infection model was similar with that of RSV A2 isolate described previously [19]. In contrast, the appearance of small clusters of infected cells was detected in rg224RSV-ΔG-infected cell monolayers by 2 and 3 dpi, and by 4 dpi intensely anti-F-stained smaller cell clusters were detected. The smaller clusters of infected cells indicated that the rg224RSV-ΔG virus exhibited impaired spread in the cell monolayer. The presence of anti-F stained debris over the cell monolayer was also observed by 3 dpi and increased by 4 dpi. This was not apparent during the early stages of infection and is consistent with localised cell damage caused by virus replication in these smaller infected cell clusters. However, this cell debris was non-infectious since it did not appear to give rise to infection in the cells that it came into contact with, even up to 6 dpi (Huong et al, unpublished observations). These data are consistent with reports that RSV lacking the G protein is impaired in HEp2 cells, and is over attenuated in animal cell models of infection [32]. In a parallel analysis, the HEp2 cell monolayers infected with rg224RSV or rg224RSV-ΔG were harvested at between 1 and 4 dpi and examined by immunoblotting using anti-G, anti-M2-1 and anti-F (Fig. 7B), which confirmed the absence of the G protein in rg224RSV-ΔG-infected cells. Although a small reduction in the F protein was observed in rg224RSV-ΔG-infected cells (presumably due to the reduced virus spread), the absence of the G protein appeared to have no general inhibitory effect on the expression of other virus genes and was consistent with previous observations (e.g. [17]).

**Figure 7.**
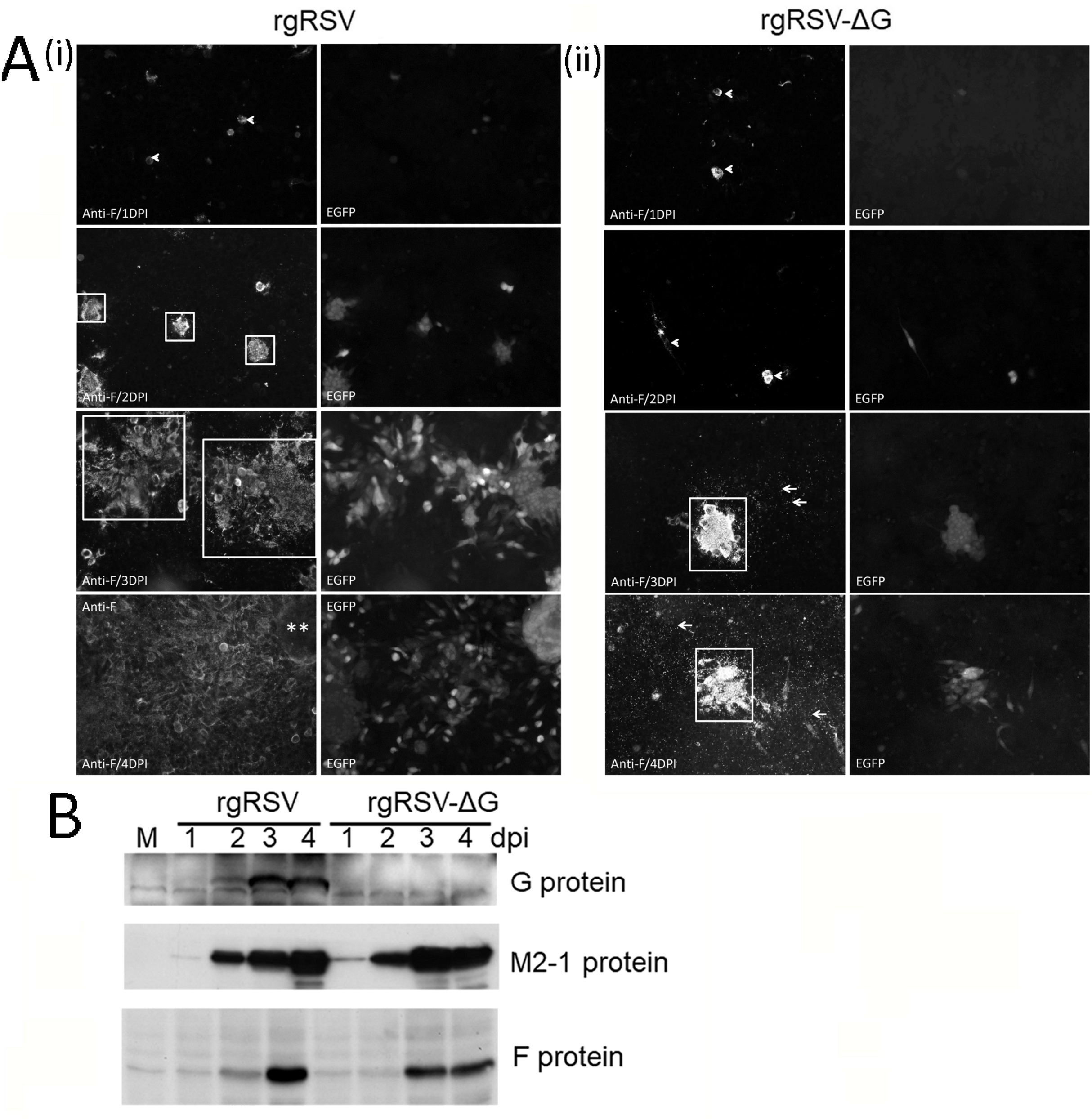
The presence of the G protein is required for efficient localised virus spread in low multiplicity of infection model. **(A)** HEp2 cells were infected with either (i) rg224RSV or (ii) rg224RSV-ΔG using a multiplicity of infection of 0.0001 and at between 1 and 4 days post-infection (dpi) the cells were stained with anti-F. The anti-F and the green fluorescent protein (EGFP) staining patterns were imaged using immunofluorescence microscopy (objective x20 magnification). The infected cell clusters are highlighted (open white boxes), the single infected cells (arrow heads) and cell debris (longer arrows) are indicated. **(B)** At between 1 and 4 dpi the rg224RSV- and rg224RSV-ΔG-infected cell monolayers were extracted in boiling mix and examined by immunoblotting using anti-G, anti-M2-1 and anti-F. Protein bands corresponding in size to the respective virus proteins are indicated. M is mock-infected.

Virus filaments have been shown to mediate virus infection and they play a role in localised cell-to-cell virus transmission [2, 3, 33], and we therefore hypothesised that the reduced virus spread observed in rg224RSV-ΔG-infected cells could be due to impaired virus filament formation. The anti-F-stained cells were therefore examined by IF microscopy at 24 hpi (Fig. 8A) and by confocal microscopy at 3dpi (Fig. 8B) to determine if virus filament formation occurred in the absence of the G protein. On rg224RSV-infected and rg224RSV-ΔG-infected cells the presence of anti-F-stained virus filaments was detected. However, to visualise the virus filaments more clearly on rg224RSV-ΔG-infected cells, the sensitivity on the camera setting had to be reduced due to the apparent increase in anti-F staining intensity on these cells. This apparent increase in antibody staining intensity was consistent with accumulation of the F protein within the smaller infected cell clusters. This analysis indicated that while virus spread was mediated less efficiently in the absence of the G protein, the absence of the G protein did appear to not inhibit the trafficking of the F protein to the site of virus assembly. Therefore, we can conclude that a block in virus filament formation could not account for the observed reduced transmission of rg224RSV-ΔG.

**Figure 8.**
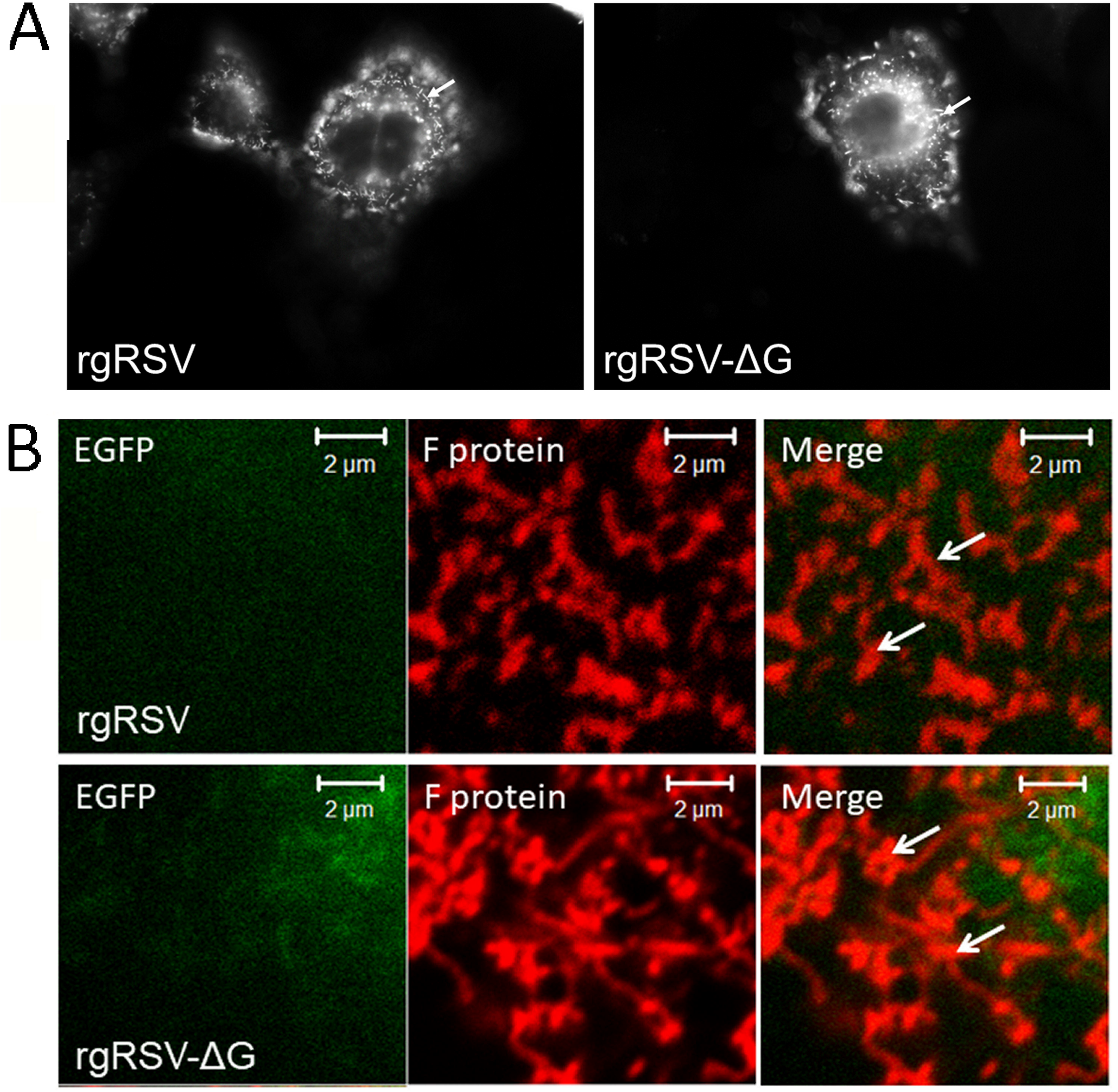
The presence of the G protein is not required for trafficking of the F protein to the sites of virus assembly. **(A)** Cells were infected using a MOI of 0.0001 and at 24 hrs post-infection the cells were stained using ant-F and examined by immunofluorescence (IF) microscopy to visualise the anti-F staining on infected cells (objective x100 (oil) magnification). **(B)** At 3 dpi the anti-F stained rg224RSV or rg224RSV-ΔG infected cells were examined by confocal microscopy to detect the presence of virus filaments. The image of rg224RSV-ΔG-infected cells was recorded using a reduced camera setting that allows the virus filaments to be more clearly imaged. The virus filaments are highlighted (white arrows).

## Conclusion

While our data suggests that the G protein was required for initiating a multiple cycle RSV infection, it does not appear to be required to infect individual cells in the cell monolayer. In this context our data is consistent with these previous report that have examined RSV isolates that do not express the G protein (e.g. [32] [17]). However, these reports have suggested that the G protein was dispensable for RSV to initiate infection in some permissive cell types. The apparent contradiction in relation to the established role of the G protein in mediating cell attachment was subsequently resolved when it was demonstrated that in these cells the F protein can also bind to susceptible cells independently of the G protein [34, 35]. In this context, low levels of localised transmission did occur in the low MOI infection model using rg224RSV-ΔG, however this process was greatly facilitated by the presence of the G protein. The expression of the recombinant RSV F protein alone is sufficient to mediate syncytial formation [36, 37], suggesting that the association of the RSV G protein with the F protein was not required to mediate membrane fusion. However, even in viruses where a functional interaction between the F and respective attachment protein has been demonstrated, the expression of the recombinant fusion protein alone can initiate membrane fusion [38]. It is therefore possible that the association of the G protein with the F protein may have other consequences. Receptor-mediated fusion involving a functional association between the respective attachment protein and virus fusion protein is a feature of many members of the paramyxoviruses [13]. In this paradigm the interaction between the attachment protein and the cell receptor leads to structural changes in the conformation of the F protein that initiates membrane fusion. In some cases both functional and structural interactions between the virus attachment and fusion proteins have been demonstrated [39–41]. During the process of virus particle assembly the F protein must be prevented from mediating the premature fusion of the virus envelope and cell membranes, as this would be expected to lead to an inhibition in virus infectivity i.e. virus inactivation. It is therefore critical that the F protein is maintained in a pre-fusion form prior to cell membrane insertion during virus entry. There are likely to be several factors that stabilise the F protein in its prefusion conformation within the virus envelope. Based on our analysis, we can speculate that the interaction between the F and G proteins may contribute to F protein stabilisation within the virus envelope, and this may be a factor in preventing premature membrane fusion. This suggestion is compatible with the results of the low MOI infection model, since although premature membrane fusion would not be expected to inhibit virus gene expression in infected cells, it would be expected to impair the localised spread of virus infection within the cell monolayer. Interestingly, the structure of the F protein that was obtained by co-crystallisation with an antibody that stabilized the F protein in its pre-fusion form [42] has demonstrated that interactions with other proteins can potentially stabilise the pre-fusion form of the RSV F protein. Furthermore the recombinant expressed F protein can be engineered to form a stable pre-fusion structure with improved and enhanced immunological properties [43, 44]. The interaction between the F and G proteins has also been demonstrated in RSV viruslike particles (VLPs) [30], suggesting that the formation of a protein complex is an intrinsic biological property of these proteins and is not dependant on virus infection. Furthermore, the VLPs containing both the F and G proteins exhibit enhanced neutralising antibody titres compared with VLPs that only contain the F protein, suggesting that the G protein may stabilise the F protein in its pre-fusion form [45]. In the context of the infectious virus, we can hypothesise that in the virus envelope the interaction with the G protein may lead to stabilization of the F protein in its pre-fusion form. Although a better understanding of the interaction between the F and G protein should lead to an improved understanding of the mechanism of virus assembly, this information may also facilitate the development of viable antiviral strategies that can destabilise this interaction. Furthermore, the interaction of the G and F proteins may have implications in the development of new RSV vaccine candidates. The presence of the G protein in such formulations could both generate antibody responses that block cell attachment [46] as well improve antibody responses against the F protein by stabilising the F protein in its pre-fusion conformation. In this context, future work will be needed to examine further the functional significance of the interaction between the F and G proteins in the virus envelope, and to further define the structural basis for the formation of this protein complex.

## Declarations

### Consent for publication

All authors approved publication

### Author contributions

TNH performed all experiments described in the manuscript, TBH and RJS wrote the manuscript. RJS conceived of the study

## Acknowledgements

We thank Mark Peebles (Nationwide Children’s Hospital in Columbus, Ohio) and Peter Collins (National Institute of Allergy and Infectious Diseases) for providing the g224RSV-FSG and g224RSV-FS viruses. We thank the Ministry of Education, Republic of Singapore (RG59/12) for providing funding (awarded to RJS).

